# PanBGC: A Pangenome-inspired framework for comparative analysis of biosynthetic gene clusters

**DOI:** 10.1101/2025.08.11.669102

**Authors:** Davide Paccagnella, Caner Bagci, Athina Gavriilidou, Nadine Ziemert

**Author notes:** Corresponding author Nadine Ziemert.

## Abstract

Bacterial secondary metabolites are a major source of therapeutics and play key roles in microbial ecology. These compounds are encoded by biosynthetic gene clusters (BGCs), which show extensive genetic diversity across microbial genomes. While recent advances have enabled clustering of BGCs into gene cluster families (GCFs), there is still a lack of frameworks for systematically analysing their internal diversity at a population scale. Here, we introduce **PanBGC**, a pangenome-inspired framework that treats each GCF as a population of related BGCs. This enables classification of biosynthetic genes into core, accessory, and unique categories and provides openness metrics to quantify compositional diversity. Applied to over 250 000 BGCs from more than 35 000 genomes, PanBGC maps biosynthetic diversity of more than 80 000 GCFs. To facilitate exploration, we present **PanBGC-DB** (https://panbgc-db.cs.uni-tuebingen.de), an interactive web platform for comparative BGC analysis. PanBGC-DB offers gene- and domain-level visualizations, phylogenetic tools, openness metrics, and custom query integration. Together, PanBGC and PanBGC-DB provide a scalable framework for exploring biosynthetic gene clusters at population resolution and for contextualizing newly discovered BGCs within the global landscape of secondary metabolism.

## Introduction

Microbial genomes are remarkably dynamic, shaped by an ongoing interplay of gene acquisition, loss, duplication, and rearrangement[1–4]. This evolutionary fluidity allows microorganisms to adapt to diverse ecological niches, develop resistance mechanisms, and expand their metabolic capacities[5–7]. As the volume of sequenced genomes continues to grow, comparative genomics has become a key approach for uncovering patterns of gene conservation and variation across related strains[8, 9].

The shift from analysing single genomes to comparing entire groups has given rise to powerful frameworks that help organize and interpret genomic diversity[10]. Among them, the pangenome model[11, 12] offers a structured view of how genes are distributed across populations, highlighting both shared and variable features that may underlie functional and ecological differences [13, 14].

The pangenome framework formalizes genomic diversity by organizing all genes found across a group of related organisms into three categories: core genes (present in all strains), accessory genes (shared by some but not all), and unique genes (found in only one strain)[9, 15–17]. This model captures both the conserved backbone of a species and its flexible genomic reservoir, providing a foundation for understanding how populations adapt, specialize, and diversify over time[13, 18, 19].

Beyond categorization, pangenomic analysis enables broader questions about genomic plasticity[19]. One such concept is pangenome openness, which quantifies how much new genetic material continues to be discovered as additional genomes are sampled[9, 18]. In open pangenomes, gene content continues to grow with each new genome, suggesting high rates of horizontal gene transfer and ecological versatility. Conversely, closed pangenomes saturate quickly, indicating more stable, conserved genetic repertoires.[17, 20] These metrics offer crucial insight into the evolutionary dynamics and adaptive strategies of microbial populations.

Over the past decade, this framework has been widely adopted in microbial genomics[21], helping to characterize evolutionary dynamics and adaptive potential in species ranging from pathogens to environmental isolates[8, 19, 22].

In recent years, similar approaches are now being extended to the study of secondary metabolism[25]. These specialized metabolites, which play key roles in microbial interactions and biotechnological applications, are encoded by biosynthetic gene clusters (BGCs)[26] (Fig.1a). BGCs exhibit complex evolutionary histories and can be considered evolutionary entities in their own right[27]. Recent studies of individual gene cluster families (GCFs), groups of BGCs that share similar biosynthetic architectures and typically produce structurally related compounds, have shown that BGCs evolve through gene gain, loss, and rearrangement, mirroring the dynamics seen in microbial genomes[28–30].

Advances in tools such as BiG-SCAPE[24] and BiG-SLiCE[23] now make it possible to group BGCs into gene cluster families and study variation within them. This enables comparative analyses of modular organization and functional divergence[24]. Yet, while such studies have begun to uncover the evolutionary dynamics within individual GCFs[31], a global, population-level framework for analysing BGC diversity has been lacking.

Here, we introduce the PanBGC framework (Fig. 1b), which applies pangenome principles to biosynthetic gene clusters by treating each GCF as a population of related clusters. This enables the systematic classification of biosynthetic genes into core, accessory, and unique components, and allows us to quantify patterns of modularity and openness across thousands of families. Applied to over 80 000 GCFs derived from more than 250 000 BGCs, the present analysis reveals that biosynthetic innovation is primarily driven by combinatorial rearrangement of conserved gene sets.

**Figure 1:**
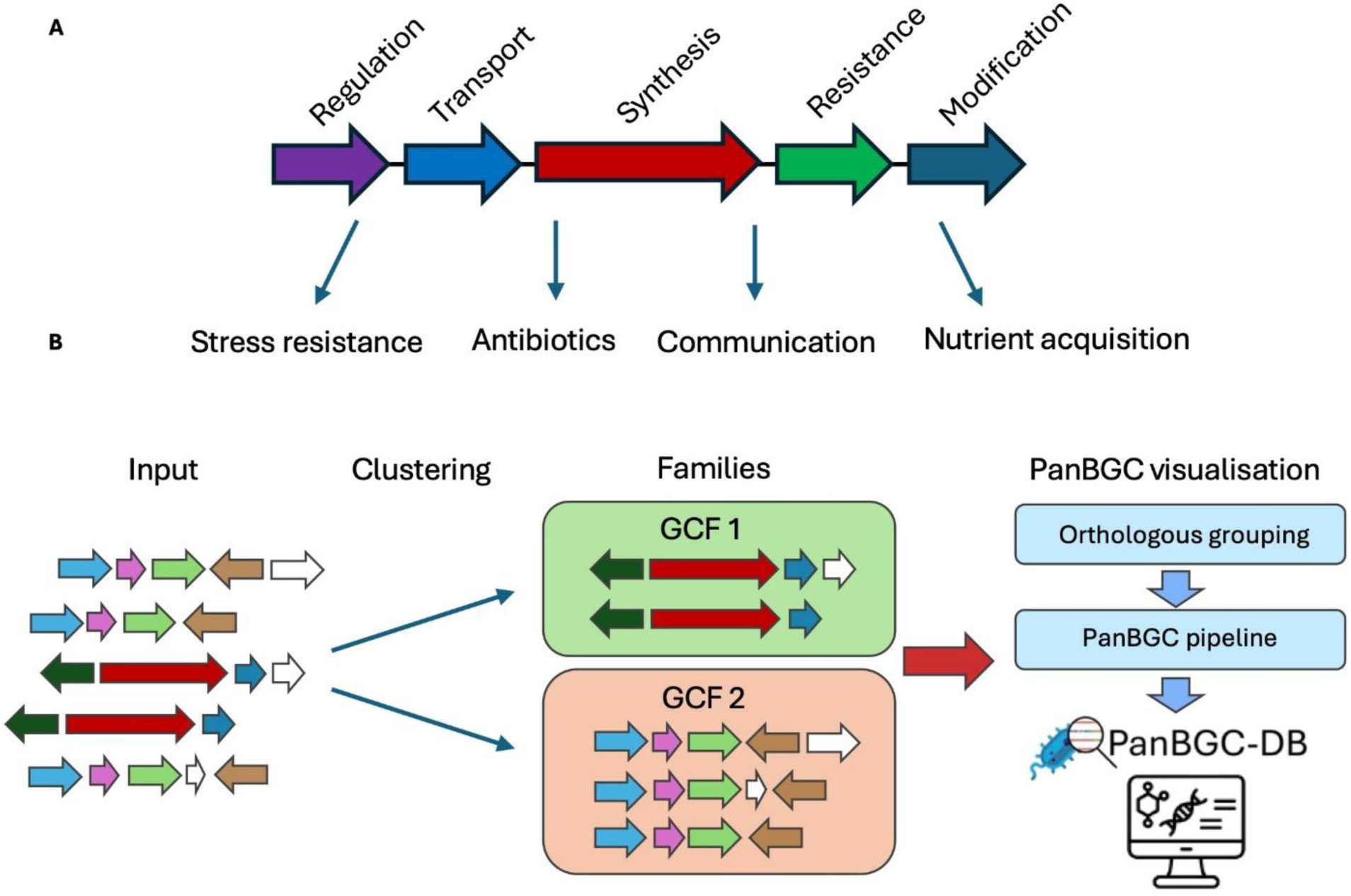
Overview of biosynthetic gene cluster (BGC) function and PanBGC-DB workflow. **a** Schematic representation of a typical BGC, highlighting the modular architecture with genes performing distinct biosynthetic functions. Each coloured arrow represents one gene in the BGC. The specialized metabolites produced by these BGCs serve diverse ecological functions in nature, like stress resistance, antibiotic production, intercellular communication, and nutrient acquisition. **b** Workflow of the PanBGC-DB pipeline. BGCs are first provided as input and clustered into gene cluster families (GCFs) based on sequence similarity and synteny by BiG-SLICE[23] and BiG-SCAPE[24]. Within each GCF, genes are grouped into orthologous groups to enable comparative analysis of core and accessory components. The resulting data are integrated into the PanBGC-DB platform for visualization and downstream exploration of BGC diversity and evolution.

To support broad access and interactive analysis, we present PanBGC-DB (https://panbgc-db.cs.uni-tuebingen.de), an open platform that enables global exploration of biosynthetic gene cluster families analysed using the PanBGC framework and integration of user-provided BGCs into this comparative landscape.

## Methods

Biosynthetic gene cluster data were compiled from two publicly available sources. A total of 254 792 predicted BGCs were obtained from the antiSMASH-DB[32] (accessed date February 2025) by automated web crawling of all available JSON-formatted annotations. In addition, we included 2 635 experimentally validated BGCs from the MIBiG v4.0 database[33], using GenBank-format files. These datasets were used as the input for clustering the BGC in gene cluster families.

### Clustering of Gene Clusters into GCFs

Biosynthetic gene clusters were clustered using a two-step strategy to define gene cluster families. First, all BGCs were processed using BiG-SLiCE[23] (v2.0.0; database release 2022-11-30) with default parameters and cut-off set to 0.7 to assign clusters to broadly defined gene cluster families based on global similarity. The computation was performed using 96 CPU cores. In the second step, each BiG-SLiCE family was subjected to fine-scale reclustering using BiG-SCAPE (v2.0-beta6)[24], which computes pairwise similarities based on domain architecture, composition and sequence similarity. BiG-SCAPE was executed with the following parameters: --input-mode recursive, --record-type region, --classify category, --gcf-cutoffs 0.4, --include-singletons, --hybrids-off, --no-trees, and --force-gbk, using the Pfam-A HMM database for domain annotation. This two-step approach allowed us to generate GCFs that reflect both broad biosynthetic relatedness and fine-grained architectural similarity.

### Orthologous grouping of genes within Gene Cluster Families

For each GCF, genes from all associated BGCs were grouped into orthologous groups using ZOL v1.5.9[34], with the -r option to standardize locus tags. Functional annotations were assigned using ZOL’s built-in annotation libraries. In addition to the ortholog assignments, we also extracted the consensus order, diversity, and average length of each orthologous group as computed by ZOL. The complete output was converted into a structured JSON format for downstream analysis and visualization.

### Openness metrics calculation

To assess openness of the gene pool, a methodology of previous research[8, 35, 36], and was adapted for the PanBGC-DB. The code is available on the GitHub repo (https://github.com/ZiemertLab/PanBGC-DB/tree/master/Website_code/public/cluster_charts/scripts) to simulate gene accumulation across each GCF. For each family, 30 random BGC sampling permutations were generated to compute the cumulative number of genes as BGCs were incrementally added. The resulting gene accumulation curves were fitted to the Heaps’ Law model y = k * x^γ^ using three approaches:

1. Standard log-log linear regression, which applies linear regression on log-transformed gene and BGC counts.
2. Weighted regression, which emphasizes later sampling points by assigning increasing weights, improving fit when early data is noisy.
3. Non-linear optimization, which directly minimizes squared error using gradient descent without log transformation.

The best-fitting model was selected based on the highest R² value. The final γ value was used to quantify the openness of each GCF.

For the analysis based on unique gene composition in a BGC, the same simulation strategy and model-fitting approaches were applied. Instead of tracking the cumulative total of all genes, only the appearance of a BGC with a unique gene composition (not including gene order) at each sampling step was counted. The resulting curves capture the rate at which novel genes appear as more BGCs are added. As before, the Heaps’ Law model was fit using all three approaches, and the model with the best R² was selected to report the final γ and k values.

To evaluate whether openness estimates based on unique gene composition differed significantly from those based on total gene count, we applied a Kruskal–Wallis rank-sum test on the corresponding γ values across all GCFs. This non-parametric test was used to assess differences in distributions without assuming normality.

### Phylogenetic tree construction

Gene trees for each orthologous group were generated by ZOL. The BGC tree is a coalescent tree inferred by astral-pro3 version 1.19.3.5[37] using all OG trees of the respective GCF.

### Website visualization

The interactive web platform was developed using JavaScript for both frontend and backend logic, with HTML and CSS for structure and styling. Visualizations were implemented using different libraries. The input data is provided in JSON, Nexus, and CSV formats (Supplementary Tab. 1). Pre-processed data is dynamically loaded and rendered client-side. Code for the website is available in the GitHub repo (https://github.com/ZiemertLab/PanBGC-DB/tree/master/Website_code/).

### Querying with user-provided BGCs

Users can upload a single BGC in GenBank format to identify the most similar GCF. For each GCF, a theoretical maximum BGC was constructed by merging all orthologous gene groups observed across its member clusters (https://github.com/ZiemertLab/PanBGC-DB/blob/master/Max_BGC.py). These representative BGCs were compiled into a searchable database using the makedb module from cblaster v1.3.0[38]. Upon upload, the user’s BGC is queried against this database using the cblaster search function to determine the best-matching GCF (Supplementary Fig.3).

### Visualization of user-generated GCFs

The Python pipeline used to process precomputed GCFs on the website was adapted to support user-provided data. Users can run this pipeline locally on one or more of their own GCFs to generate a structured JSON file compatible with the platform. By uploading this file, users can visualize their GCFs using the same interface and features as the preloaded dataset. The pipeline is available under https://panbgc-db.cs.uni-tuebingen.de/data/Scripts.zip.

## Results

Building on the conceptual framework introduced before, we adapted the pangenome model to the analysis of BGCs. In this context, each gene cluster family is treated as a population-level unit analogous to a microbial species. This analogy is grounded in the fact that BGCs grouped into the same GCF share high architectural and functional similarity[39], often producing structurally related but still distinct compounds. Like individual genomes within a species, the BGCs within a GCF represent naturally occurring variants that reflect evolutionary diversification around a conserved biosynthetic theme[29]. This enables the application of core, accessory, and unique gene classification to BGCs, allowing us to investigate their diversity not as isolated cases but as structured populations with internal variability.

### Construction and Clustering of the PanBGC-DB Dataset

To build a comprehensive dataset capturing bacterial BGC diversity, data from two major resources was compiled: the antiSMASH database[40], providing 254 792 BGCs from 35 726 bacterial genomes, and the MIBiG repository[33], contributing an additional 2 635 experimentally characterized BGCs.

To generate refined GCFs a two-stage clustering strategy was used. Initial clustering using BiG-SLiCE grouped 257 427 BGCs into 21 528 GCFs. Notably, 15 443 GCFs consisted of singletons (BGCs that did not cluster with any other BGC), representing ∼6% of the total BGC dataset. To further refine family delineation each GCF was subsequently clustered using BiG-SCAPE v2.0, yielding a final set of 80 698 unique families. Of these, 58 700 GCFs were singletons, representing ∼23% of the total BGC dataset. This indicates that the second clustering step identified additional fine-scale distinctions among loosely related BGCs (Fig.2a).

Excluding singleton families, the average size of the refined GCFs was 9.4 BGCs (Supplementary Fig.1). These BGCs spanned 58 classes including nonribosomal peptide synthetases (NRPS), polyketide synthases (PKS), terpenes, ribosomally synthesized and post-translationally modified peptides (RiPPs), saccharides, others and diverse hybrid combinations thereof. Non-hybrids BGCs have the most GCFs, while hybrids of three or more different classes build the least GCFs (Fig. 2b)

**Figure 2:**
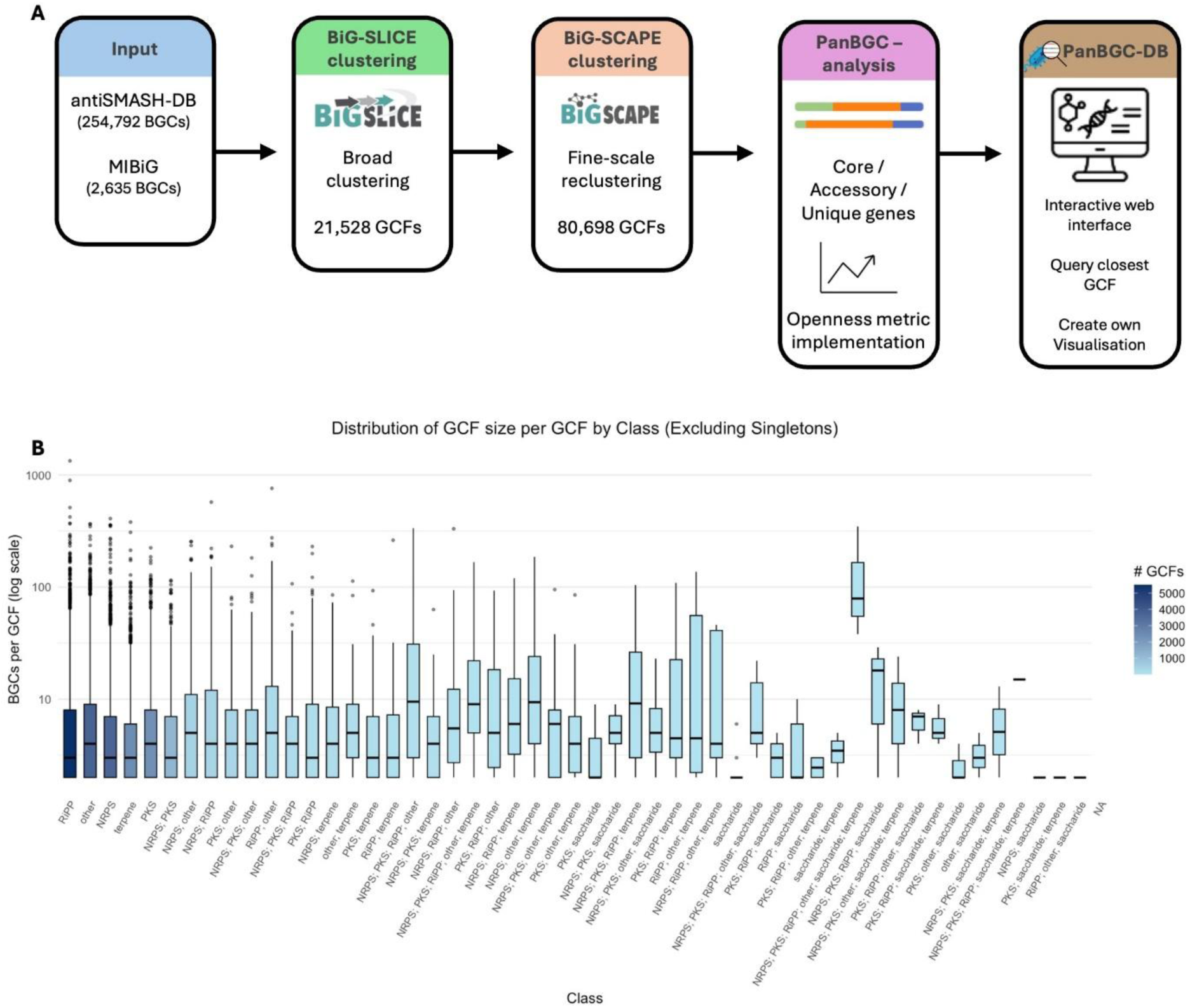
PanBGC-DB pipeline overview and BGC family size distribution. **a** BGCs from antiSMASH-DB and MIBiG are first clustered using BiG-SLiCE. Broad GCFs are further refined using BiG-SCAPE. The PanBGC analysis framework then identifies core, accessory, and unique genes within each family and implements openness metrics. The results are integrated into the PanBGC-DB platform, which provides an interactive web interface for querying, visualization, and user-uploaded comparisons. **b** Distribution of GCF sizes across classes, excluding singletons. Each boxplot shows the number of BGCs per GCF (log scale) for a given class, with colour gradient indicating the number of GCFs per class.

### Core and Accessory Gene Classification within GCFs

Following standard pangenomic practice[8, 21, 41], all genes from BGCs belonging to the same GCF were clustered into orthologous gene groups (OGs). Each OG was subsequently classified as core, accessory, or unique according to its frequency across the clusters in that family. In bacterial pangenome studies, genes present in at least 95 % of genomes are typically considered core[12, 20, 21]. Because BGCs contain far fewer genes, applying a 95 % cutoff risks inflating the core fraction when only a handful of clusters differ and so loosing valuable information about GCF diversity. To avoid this bias, we adopted a more stringent criterion: an OG was deemed core only if it occurred in 100 % of BGCs within the family. OGs present in more than one but not all clusters were classified as accessory, whereas OGs exclusive to a single BGC were labelled unique. This stricter definition preserves meaningful distinctions between conserved and variable functions and provides a clearer view of genetic diversity within each GCF.

### Functional Domain-Level Analysis

To explore trends in the functional roles of genes within BGCs, we analysed the Pfam domain annotations[42] associated with core, accessory, and unique orthologous groups across biosynthetic categories. Core genes were predominantly linked to essential enzymatic activities required for metabolite biosynthesis. For example, in NRPS clusters, condensation (C) domains were consistently classified as core due to their universal role in peptide bond formation[43]. Similarly, KS (ketosynthase) domains were frequently core in PKS systems[44]. In addition to biosynthetic enzymes, transporter-related domains were also commonly identified among core genes, reflecting the importance of compound export in BGC function.

Accessory genes displayed greater functional variability and were often associated with tailoring reactions or regulatory roles. By examining which accessory genes are recurrently present across BGCs within the same biosynthetic class, this analysis also highlights conserved auxiliary functions that may contribute to structural diversification or pathway regulation.

A complete overview of domain frequency distributions for core, accessory, and unique genes across all biosynthetic categories is available at https://panbgc-db.cs.uni-tuebingen.de/stats under the gene stats tab.

### Compositional Insights and Boundary Considerations in Intra-Family BGC Comparisons

One of the key advantages of applying a pangenomic framework to BGCs is the ability to systematically compare gene composition within a gene cluster family and link these differences to chemical diversity. For instance, in a GCF containing experimentally validated Malleobactin A–D[45–48] and Ornibactin C4–C8 clusters[49], we identified clear substructuring based on accessory genes that correlate with distinct chemical features. Malleobactin-producing BGCs encode a formyltransferase absent from Ornibactin clusters, while the Ornibactin subset consistently features two acyltransferase genes not found in Malleobactin variants (Fig. 3a). These accessory elements are mutually exclusive and align with known structural modifications in their respective siderophores (Fig. 3b-c), highlighting the utility of the framework in pinpointing biosynthetic genes responsible for functional diversification. This comparative resolution also opens up possibilities for synthetic biology applications, where distinct accessory genes can be rationally combined to design novel hybrid clusters with tailored chemical output. Expanding from this example, systematic identification of gene-function relationships across thousands of GCFs paves the way for automated prediction of accessory gene functions and chemical modifications based on gene content, potentially accelerating the discovery and engineering of novel bioactive compounds.

**Figure 3:**
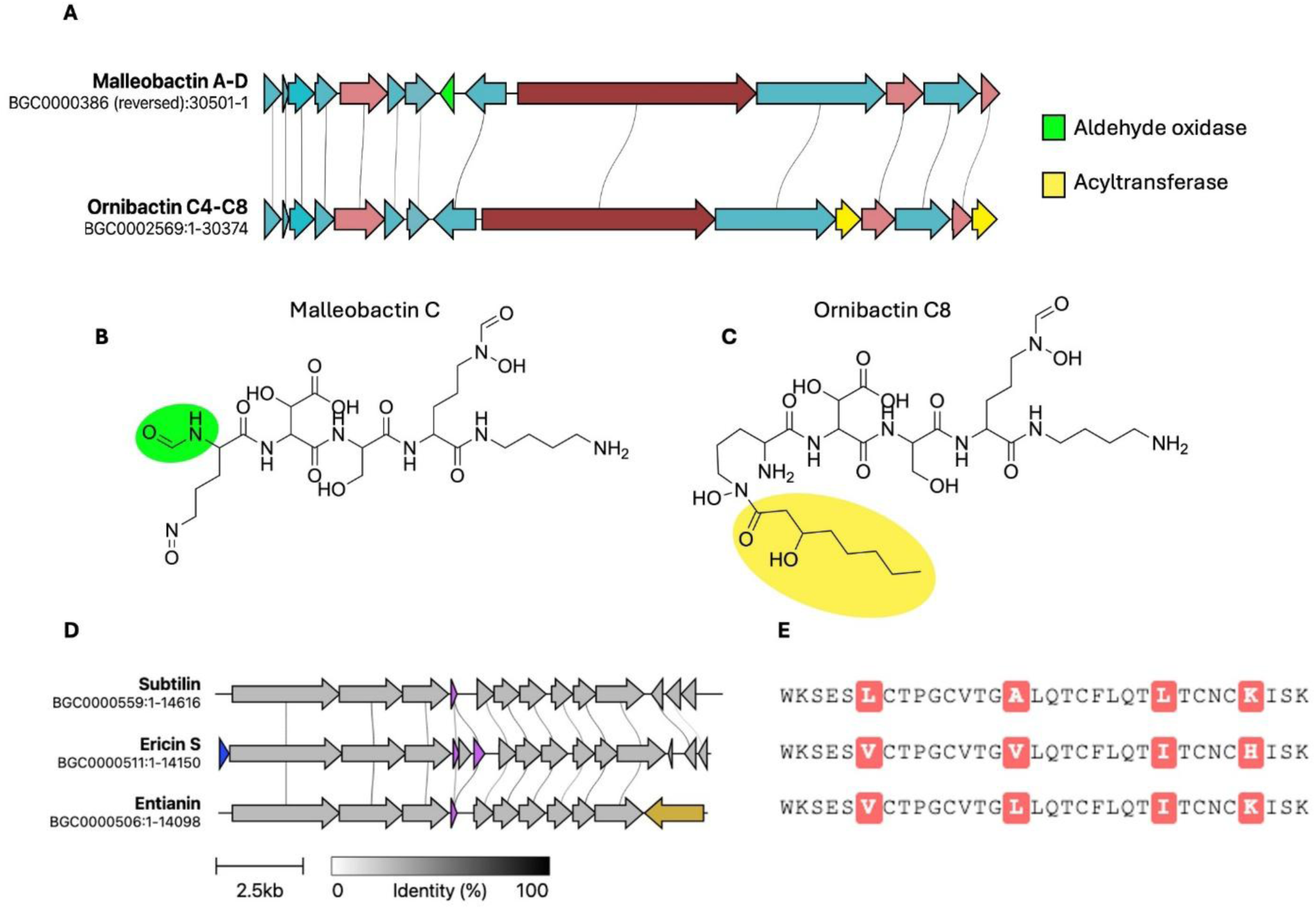
Gene cluster variation and structural diversification in related siderophores. **a** Comparative gene cluster alignment of the Malleobactin A–D and Ornibactin C4–C8 BGCs from the MIBiG database. Homologous genes are connected by grey lines. While both clusters share a conserved core biosynthetic architecture, they differ in accessory genes: the Malleobactin cluster encodes an aldehyde oxidase (green), whereas the Ornibactin cluster features additional acyltransferases (yellow). **b,c** Chemical structures of Malleobactin C and Ornibactin C8. Structural differences introduced by the respective accessory genes are highlighted with the respective colors. The remaining structure is shared by both compounds. **d** Compositional differences of the Subtilin, Ericin S and Entianin BGC. In blue a unique Ericin S transporter, in purple the duplicated structural genes and in brown the unique regulator of Entianin. **e** Core peptide alignment of Subtilin, Ericin S and Entianin.

However, interpreting such compositional differences also requires caution due to limitations inherent in automated boundary predictions. As the BGC definitions in our dataset rely on antiSMASH outputs, the predicted cluster boundaries are often influenced by synteny and genomic context.[32] This can lead to the inclusion of conserved flanking regions that are not functionally related to the BGC, particularly in closely related species where genomic neighbourhoods are similar. This challenge is exemplified by the GCF containing the enterobactin pathway. The MIBiG entry for enterobactin (BGC0002685) and amonabactin P 750 (BGC0001502) define compact and experimentally validated clusters[50, 51], yet related BGCs detected from antiSMASH extend considerably beyond this boundary, capturing additional genes on both ends (Fig.4). It is unclear whether these genes represent novel accessory functions or unrelated genomic content.

**Figure 4:**
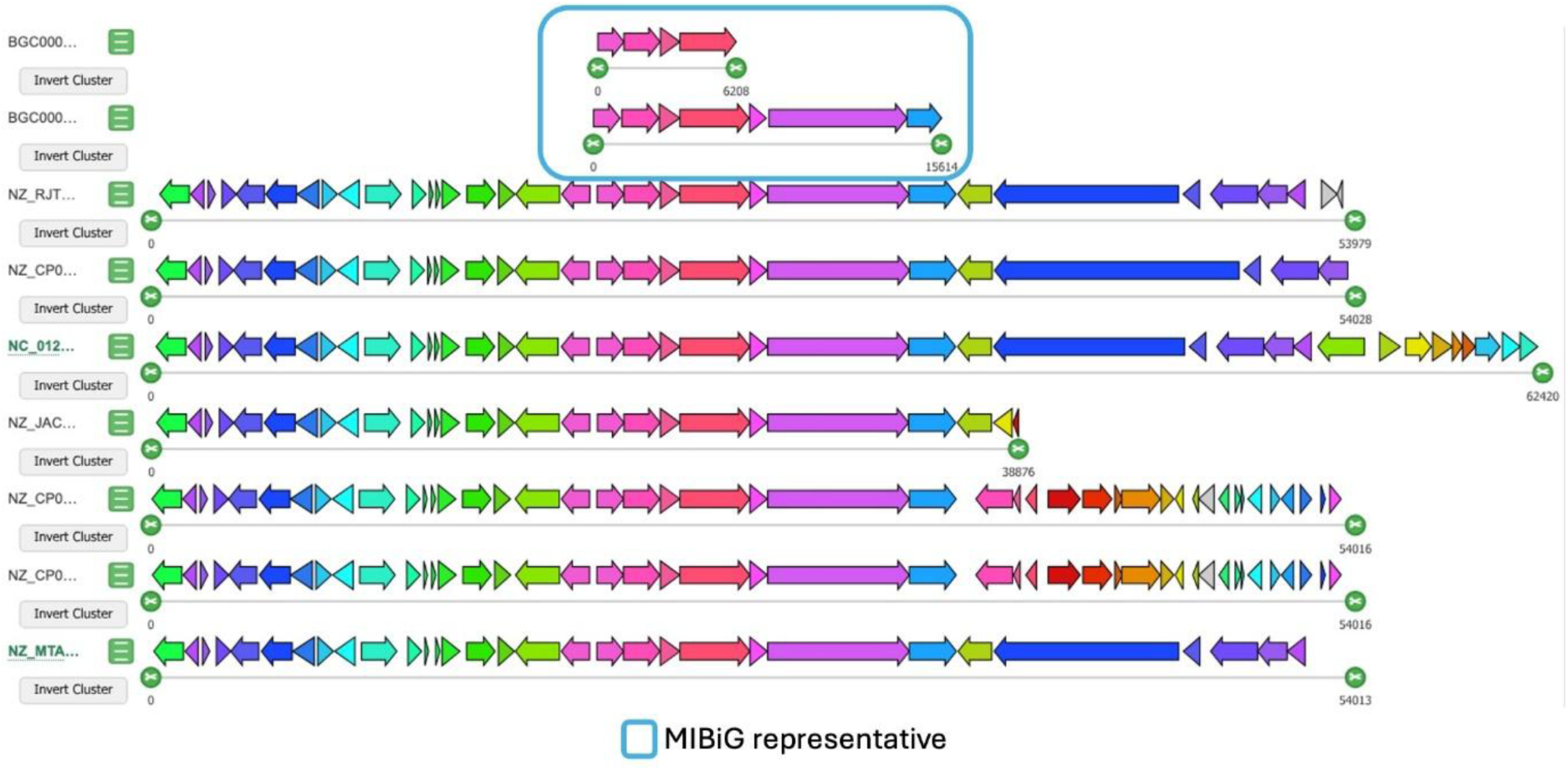
Gene cluster diversity within the Enterobactin and amonabactin P 750 family. Comparative visualization of BGCs belonging to the Enterobactin gene cluster families. Each row represents a single BGC, with genes colored by orthologous group and arrows indicating gene orientation. MIBiG reference cluster is highlighted with a blue box.

Despite this limitation, the comparative approach offers a strategy to refine cluster boundaries. When antiSMASH-predicted clusters can be directly compared with a curated MIBiG entry, shared core regions can be confidently delineated. Accessory genes that are consistently absent in MIBiG but variably present in antiSMASH predictions may be deprioritized for functional interpretation. In this way, the pangenome model not only supports the discovery of biosynthetic variability but also provides a framework for improving cluster annotation fidelity through comparative context.

### Gene Composition Openness Reveals Modular Innovation in BGCs

To quantify the degree of compositional variability within gene cluster families, we adapted openness metrics from microbial pangenome analysis using Heaps’ law γ-values [8, 12]. These metrics assess whether the gene content within a family is relatively saturated (closed) or continues to expand with the inclusion of new members (open), reflecting either genetic stability or ongoing diversification. Unlike species-level pangenomes, where hundreds to thousands of genomes are typically analysed, most GCFs consist of only a few BGCs. To address this constraint, we implemented modified curve fitting strategies tailored to smaller sample sizes and restricted openness analysis to GCFs with at least three BGCs, as reliable γ-value estimation was not feasible for smaller families.

To capture different dimensions of diversity, openness was defined in three distinct forms. First, gene-based openness quantifies the increase in the total number of orthologous groups (OGs) with each additional BGC, reflecting expansion of the overall gene repertoire (Fig. 5a). Second, composition-based openness measures how consistently OGs are reused across BGCs in a GCF, indicating variability in how subsets of the PanBGC are deployed. This captures the rate at which novel gene combinations (distinct sets of orthologous groups) appear with each additional BGC, regardless of gene order (Fig. 5b). Third, we also considered the rate at which entirely novel genes, those not previously observed in a family, appear with the addition of new BGCs. However, our main focus remained on gene- and composition-based openness, which together capture both repertoire size and modular flexibility.

**Figure 5:**
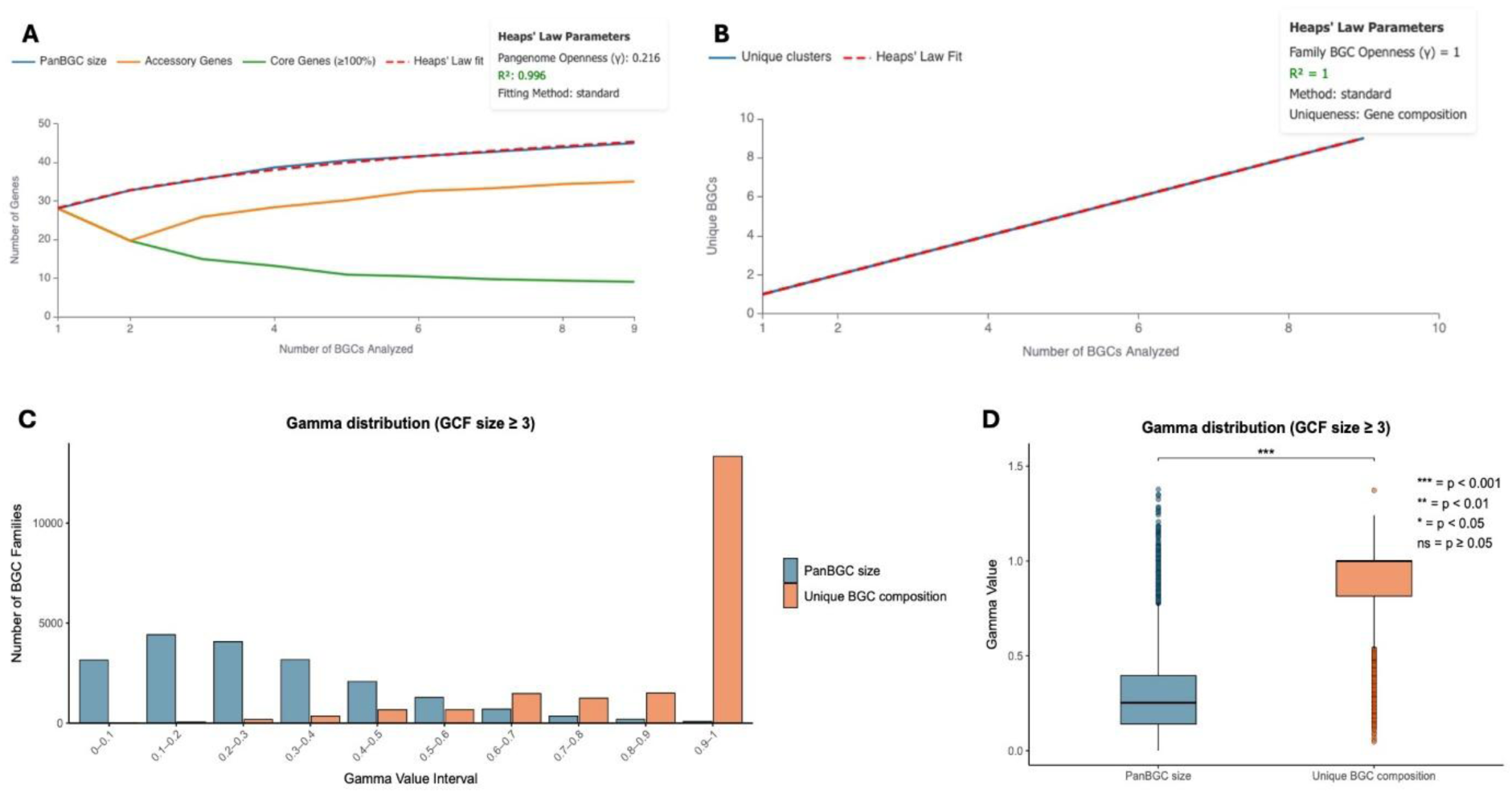
Openness metrics adapted for biosynthetic gene clusters. **a** Example Heaps’ Law curve showing the accumulation of total (blue curve), core (green curve), and accessory genes (orange curve) as additional BGCs are sampled from a representative GCF. The red dotted line represents the fitted curve. **b** Heaps’ Law modeling of BGC uniqueness using gene composition data. The red dotted line represents the fitted curve. **c** Histogram of gamma values for BGC families with ≥3 members, calculated using either PanBGC size (blue) or unique gene composition (orange). **d** Boxplot comparing gamma distributions between the PanBGC size and unique BGC composition metrics for GCFs with ≥3 BGCs n =14 716.

Among the 80 698 GCFs in the final dataset, 14 716 families contained at least three BGCs and were retained for openness calculations. Using gene-based openness (i.e., increase in PanBGC size with each added BGC), the average γ-value across all biosynthetic categories was 0.286. According to established thresholds (γ < 0.3: closed; 0.3-0.6: intermediate; >0.6: open)[8, 36], this indicates that most GCFs are closed, with relatively stable gene repertoires. This suggests a limited influx of novel genes as more BGCs belonging to the same GCF are sampled. In contrast, openness based on gene composition diversity within BGCs (i.e., how consistently subsets of PanBGC genes appear across clusters) showed an average γ-value of 0.841, indicating a high degree of structural variability (Fig. 5c). These findings highlight that gene composition reshuffling, rather than acquisition of new genes, is the dominant driver of diversity within GCFs. This suggests that in natural systems, BGC diversity emerges primarily through modular reorganization of existing genes rather than through frequent incorporation of entirely novel genes, reinforcing the idea that structural variability is a key evolutionary mechanism in BGC innovation. A Kruskal-Wallis test comparing γ-values from PanBGC size-based and composition-based openness confirmed a statistically significant difference (p < 0.001) between the two distributions (Fig.5d).

The *Bacillus* lanthipeptide family (family 415_FAM_00315) exemplifies this pattern. The clusters belonging to this family encode for subtilin, ericin S and entianin, which are antimicrobial peptides with potential applications as natural food preservatives and medicine[52]. While all three clusters share conserved core genes for lanthionine ring formation they exhibit compositional differences: the ericin cluster contains duplicated structural genes separated by an inserted *lanC* fragment, while the entianin cluster features different regulatory genes compared to the standard subtilin architecture (Fig. 3d-e)[53–55]. This family shows low gene-based openness (γ = 0.195) reflecting limited novel gene acquisition, yet a relative high composition-based openness (γ = 0.641) indicating modular rearrangement of existing genes.

To ensure that openness values were not biased by differences in family size, we also assessed correlations between γ-values and the number of BGCs per GCF (Supplementary Fig.2). No significant association was detected, confirming that openness metrics robustly reflect compositional dynamics independent of cluster count.

### Evolutionary Dynamics Assessed Through Tanglegram Analysis

To explore the evolutionary relationships among gene clusters within each family, we applied tanglegram-based comparisons between gene trees and BGC trees. Phylogenetic trees represent evolutionary relationships, with closely related sequences grouped on nearby branches. In our analysis, gene trees show the evolutionary history of individual orthologous groups (how each gene family evolved), while BGC trees represent the overall evolutionary relationships between entire gene clusters (how the complete BGCs are related to each other). For all GCFs with at least three BGCs, individual gene trees were constructed for each orthologous group and aligned against the BGC tree. These tanglegrams visually represent the congruence between gene-level and cluster-level relationships, enabling the identification of structural similarities or differences across BGCs in a family.

In some GCFs, gene trees closely mirrored the coalescent BGC tree, indicating structural consistency and shared evolutionary trajectories. In contrast, tanglegrams with extensive crossover lines suggest high plasticity of the GCF with possible horizontal gene transfers. Families with many crossover lines suggest higher evolutionary plasticity and potential for gene module exchange or recombination, while minimal crossovers indicate structurally conserved BGCs.

### Web-Based Visualization and User Tools for Exploring BGC Diversity

To make this conceptual framework accessible and interpretable, we developed PanBGC-DB (https://panbgc-db.cs.uni-tuebingen.de/), an interactive web platform for exploring biosynthetic gene cluster families (Fig. 6 a-h). The website allows users to browse thousands of precomputed GCFs (Fig. 6a) and interactively visualize their internal diversity. By presenting the results of the pangenome adaptation in an intuitive, visual format, PanBGC-DB provides a practical entry point into the population-level analysis of BGCs.

**Figure 6:**
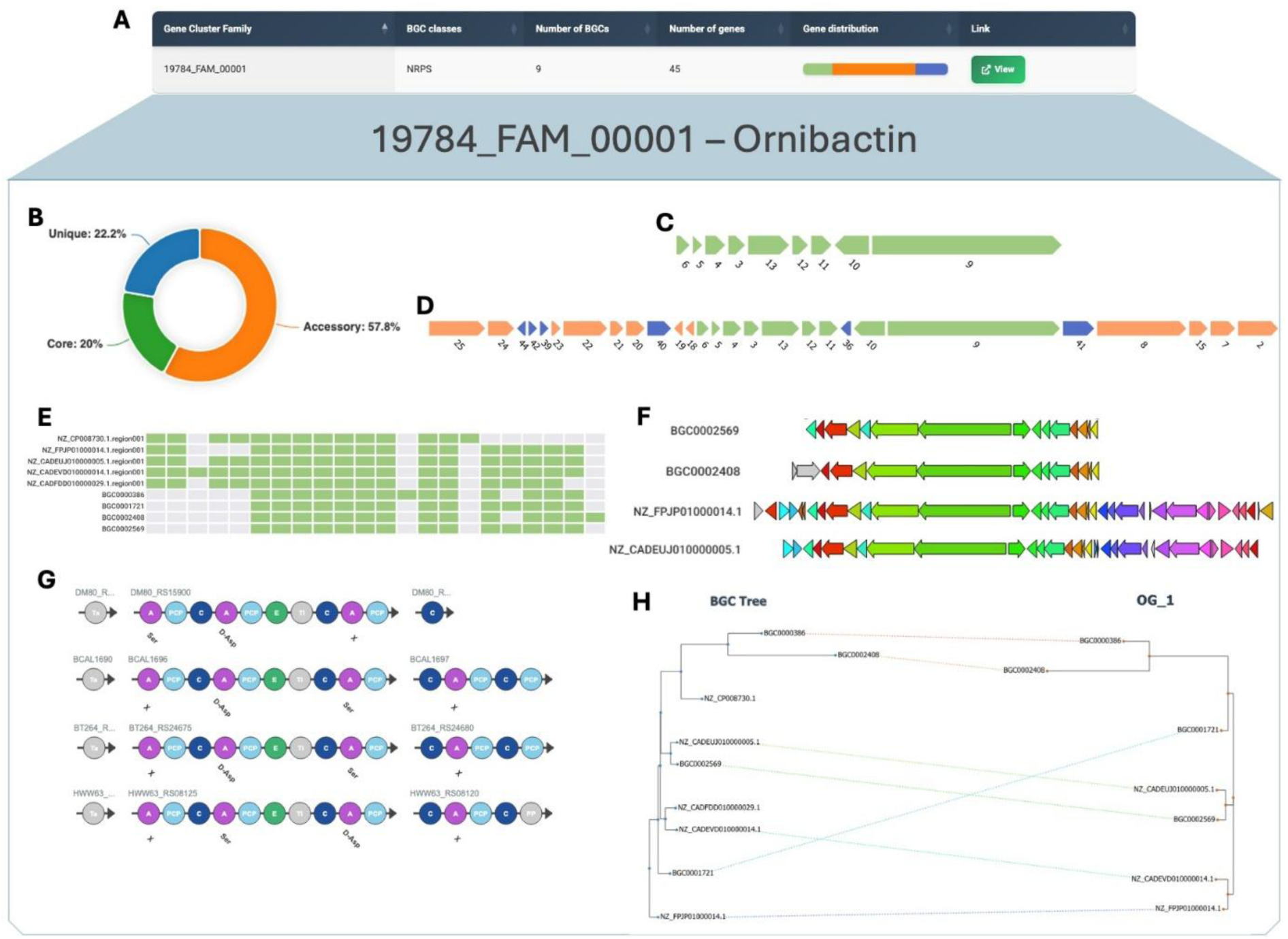
Multiple screenshots from the PanBGC-DB web interface for the Ornibactin and Ornibactin GCF. **a** Overview table showing metadata of gene cluster families, including class, number of BGCs, total genes and summary of gene distribution (core / accessory / unique). **b** Donut plot illustrating the proportion of core, accessory, and unique genes in this GCF. **c** Visual representation of the core BGC. **d** Visual representation of the maximum BGC (PanBGC). **e** Gene presence-absence heatmap across all BGCs within the family. **f** Gene cluster comparison plot showing structural conservation across BGCs. **g** Domain architecture viewer showing module organization in biosynthetic genes across the family. **h** Interactive tanglegram linking the phylogenetic tree of BGCs (left) with that of a selected orthologous group (right).

Each GCF page offers a suite of interactive modules enabling in-depth exploration of its composition and structure. Users can adjust the core gene threshold dynamically (e.g., from 100% to lower cutoffs), which updates the classification of core, accessory, and unique genes across the cluster family (Fig. 6b). Based on this threshold, the platform reconstructs a core

BGC (comprising only genes that meet the cutoff) (Fig. 6c) and a maximum BGC (all genes observed in any cluster) (Fig. 6d), both displayed in consensus gene order. These representations allow for immediate insight into conserved biosynthetic cores versus variable extensions.

To evaluate diversity within a GCF, the platform provides openness visualizations, including curves for both the increase in PanBGC size and compositional diversity as more BGCs are added (Fig. 5 a-b). These charts help interpret whether a family is genomically saturated (closed) or still expanding (open), capturing both the total repertoire and the variability in gene usage.

Further modules include a presence-absence heatmap showing the distribution of orthologous groups across BGCs in a family (Fig. 6e), and a clinker[56] inspired synteny view that groups structurally identical BGCs and aligns them for side-by-side comparison (Fig. 6f). For modular BGCs such as NRPS and PKS, domain-level annotations are visualized, allowing users to assess differences in enzymatic architecture and modular composition across clusters (Fig. 6g).

To explore evolutionary dynamics, the platform features tanglegram visualizations, comparing individual gene trees to the coalescent BGC tree (Fig. 6h). These allow users to assess evolutionary congruence or structural rearrangements within each GCF. A high number of crossover lines between trees suggests evolutionary plasticity or potential horizontal gene transfer, while low crossover indicates conserved gene arrangements.

Beyond internal visualization, PanBGC-DB also includes two tools that extend its utility to custom datasets and external queries:

#### 1. Custom BGC Visualization Pipeline

Users can download a Python-based pipeline that processes user-provided BGCs into PanBGC-style visualizations. With a single command, the pipeline generates interactive displays including core/accessory gene maps, heatmaps, domain alignments, and presence-absence matrices. This allows researchers to analyse their own gene clusters within the same conceptual framework used in the public database.

#### 2. Query Interface for Cluster Matching

A built-in search function enables users to upload a BGC of interest and identify the closest matching GCFs in the PanBGC-DB reference dataset using the cblaster tool. The query returns the most similar families based on gene content similarity, allowing users to contextualize their BGC within broader patterns of diversity and conservation (Supplementary Fig.3).

Together, these features make PanBGC-DB not only a static repository of precomputed clusters, but also a dynamic environment for hypothesis generation, comparative analysis, and integration of user-generated data within a population-level framework of BGC diversity.

## Discussion

In this study, we introduce a conceptual shift in the analysis of biosynthetic gene clusters by adapting the pangenome framework to the level of gene cluster families. Rather than treating BGCs as isolated genomic islands, we organize them into structured populations of related clusters, enabling comparative analyses grounded in evolutionary principles. This approach positions each GCF analogously to a microbial species in classical pangenomics[12, 16], allowing us to systematically partition biosynthetic diversity into core, accessory, and unique components [12, 57]. By doing so, we provide a scalable framework for interpreting the modular architecture of secondary metabolism across large genomic datasets and open the door to population-level reasoning in a domain traditionally dominated by individual-case studies. To translate this conceptual shift into an accessible resource, we developed PanBGC-DB, an interactive web platform that enables users to explore population-level diversity of BGCs across thousands of gene cluster families.

While other resources exist for organizing biosynthetic gene clusters at scale [23, 24], including the widely used BiG-FAM database [39], these platforms have primarily focused on high-level classification and dereplication of BGCs across global datasets. Workflow tools like BGCflow similarly examine BGC distribution across pangenomes, treating entire clusters as discrete genomic units[58]. In contrast, PanBGC-DB is tailored toward the in-depth analysis of variation within gene cluster families. By adopting a population-genomic framework and applying a two-step clustering strategy, PanBGC-DB generates more granular GCFs, allowing closely related BGCs to be studied as coherent evolutionary units. This finer resolution is not intended to replace broader classification schemes, but rather to support the complementary goal of revealing how modular rearrangement, gene loss, and duplication shape the diversity within biosynthetic lineages. In doing so, PanBGC-DB extends the comparative scope of BGC analysis beyond mere grouping, towards understanding the internal dynamics that drive natural product diversification.

The openness metrics for BGC gene pool and BGC composition introduced here provide a valuable framework for interpreting the evolutionary and functional potential of biosynthetic gene clusters. Our analysis revealed that while most GCFs exhibit limited acquisition of entirely novel genes, indicating a relatively closed gene content, the combinations in which these genes appear across BGCs of the same GCF are highly variable. This compositional fluidity suggests that the primary driver of BGC diversification is not the continual integration of new biosynthetic genes, but rather the modular reshuffling of a conserved gene set. This modularity is exemplified by the *Bacillus* lanthipeptide family, where conserved ring-forming enzymes are coupled with variable resistance, transport, and regulatory modules. Such flexibility allows organisms to fine-tune metabolic pathways, generate structurally distinct metabolites, and adapt to shifting ecological contexts, all without expanding their gene pool.[53, 59–61] These findings position gene composition plasticity as a central mechanism of biosynthetic innovation and underscore the value of viewing GCFs as evolutionary populations rather than isolated units. Thus, modular reshuffling allows organisms to repurpose existing biosynthetic elements into new configurations, enabling the generation of diverse metabolites without the need to acquire entirely novel genes. This strategy not only supports metabolic adaptability across ecological niches[60] but may also facilitate the emergence of novel functions by reassembling familiar parts in previously untested ways.

The ability to dissect biosynthetic gene clusters at the population level opens new opportunities for natural product discovery and design. By clearly distinguishing conserved core genes from variable accessory components within GCFs, PanBGC-DB helps researchers identify families with modular flexibility, which often correlates with chemical novelty[62]. This makes it possible to prioritize gene cluster families that exhibit unexplored biosynthetic potential, particularly those with unusual combinations of biosynthetic domains or accessory genes. At the same time, the structured view of naturally (co-) occurring genes[63] provides a valuable basis for synthetic biology, offering a blueprint for reconstructing or modifying pathways using genes present. By using gene combinations that are already observed together in nature and present in the same gene cluster family, synthetic biology can draw on pathways that are more likely to be functionally compatible. [29, 64–66] A concrete example of this design-guiding potential is illustrated by the Ornibactin/Malleobactin gene cluster family. Using PanBGC-DB, we identified accessory gene differences, specifically two acyltransferases in Ornibactin C4–C8 and a formyltransferase in Malleobactin A–D, that correlate with known structural variations between the compounds [67, 68]. These structural differences translate into functional divergence. Ornibactin C8 exhibits strong siderophore activity, while most malleobactins require concentrations exceeding 400 μM for minimal iron-chelating function. Additionally, malleobactin and Ornibactin display a different role during infection. Ornibactins are essential virulence factors for *B. cenocepacia* pathogenesis, whereas malleobactins are dispensable for *B. pseudomallei* virulence. The accessory gene modifications therefore appear to have specialized ornibactin for virulence-associated iron acquisition, while malleobactins may have evolved broader alternative biological functions. Based on this observation, we propose the rational construction of a hybrid cluster incorporating both functionalities, potentially generating a novel metabolite that combines the potent iron-chelating capacity of ornibactins with the alternative biological activities of malleobactins.[69] This demonstrates how PanBGC-DB can move beyond passive exploration to actively guide the design of new biosynthetic pathways, grounded in observed natural configurations and evolutionary compatibility. This increases the chances that genes will integrate successfully into engineered systems, both structurally and biochemically. In this way, PanBGC-DB provides a practical and biologically grounded resource for guiding pathway engineering with greater precision and confidence.

While PanBGC-DB provides a scalable framework for analysing biosynthetic gene clusters at the population level, limitations of the platform should be acknowledged. One of the key trade-offs in our approach lies in the GCF construction. By using a two-step clustering strategy to ensure that only closely related BGCs are grouped together, we enhance the resolution of within-family comparisons and make diversity patterns more interpretable. However, this increased specificity may come at the cost of excluding more distantly related clusters that, while functionally relevant, fall outside the defined similarity thresholds. As a result, broader biosynthetic relationships might be fragmented across multiple families, potentially limiting comparative insights at higher levels of divergence. In addition, the BGCs used in this study were sourced from the antiSMASH-DB, where cluster boundaries are inferred based on the position of core biosynthetic genes and domain architecture, but are not experimentally validated. As such, inaccuracies in boundary prediction may lead to the inclusion of non-functional genes or the omission of relevant tailoring enzymes, which could dilute or distort the inferred gene content of a family. Moreover, the high number of singleton clusters observed in our analysis may reflect the uniqueness of these biosynthetic systems but could also be artificially inflated by the predominance of cultivated strains in the antiSMASH database, which may not fully represent the true diversity of microbial communities and could therefore lead to an incomplete or skewed assessment of BGC diversity patterns. However, the growing interest in metagenomics and the increasing incorporation of metagenomic-derived BGCs into databases should help mitigate this bias in future. Finally, orthologous gene groups in PanBGC-DB are determined using ZOL[34]. ZOL clusters genes based on sequence similarity and positional conservation across gene clusters, which is highly scalable but may group structurally similar yet functionally divergent genes into the same OG. This can blur subtle functional differences, potentially reducing the resolution of accessory versus core gene identification. However, functionally precise orthology inference is an open problem[70], and ZOL is one of the tools performing very well with cluster genes.

Despite these limitations, PanBGC-DB represents a significant advance in the systematic exploration of biosynthetic diversity. By reframing gene cluster families as structured populations, the platform provides a powerful conceptual and analytical foundation for understanding how secondary metabolism evolves, diversifies, and adapts. Its integration of scalable clustering, gene-level pangenomic metrics, and interactive visualization tools makes the database a unique and accessible resource for both hypothesis-driven research and data exploration. As new genomes and metagenomes continue to be sequenced, and as tools for BGC boundary refinement and gene function prediction advance, the precision and utility of PanBGC-DB will continue to grow. The structured, population-level representation of BGCs provided by PanBGC also creates new opportunities for machine learning applications. For example, core/accessory gene classification enables standardized feature extraction for models predicting metabolite structure, function, or ecological role. Openness scores and domain architectures provide rich quantitative descriptors for prioritizing BGCs with biosynthetic novelty, while curated GCFs define empirically co-occurring gene sets that are valuable for training generative models. We anticipate that PanBGC will serve as a foundational resource for both experimental and computational advances in secondary metabolism.

## Data availability

PanBGC-DB is freely available at https://panbgc-db.cs.uni-tuebingen.de and can be accessed using any web browser with JavaScript support. The full source code for the website, as well as all scripts and pipelines used for clustering, orthologous grouping, and openness calculations, are available at https://github.com/ZiemertLab/PanBGC-DB.

## Acknowledgments

D.P. and N.Z. were supported by H2020-FNR-11-2020: SECRETED (grant agreement: 101000794). N.Z. was supported by the German Center for Infection Research (TTU09.717); AG was supported by the Deutsche Forschungsgemeinschaft (DFG; Project ID # 398967434-TRR 261). CB was supported by the German Center for Infection Research (TTU09.716); The authors thank the Cluster of Excellence: EXC 2124: Controlling Microbes to Fight Infection (CMFI, project ID 390838134) for the structural support. We thank the Interfaculty Institute for Biomedical Informatics (IBMI) at the University of Tübingen for providing the computational resources.

## Author contributions

D.P. and N.Z. wrote the main manuscript and designed the research; D.P. build the pipeline and created the Web interface; C.B. conducted the GCF creation; A.G. validated the tools used for orthologous clustering.

## Competing interests

The authors declare no competing interests.

## Notes

https://github.com/ZiemertLab/PanBGC-DB

https://panbgc-db.cs.uni-tuebingen.de/

### Competing Interest Statement

The authors have declared no competing interest.

