## Supplementary Figures and Table for "PanBGC: A Pangenome-inspired framework for comparative analysis of biosynthetic gene clusters"

### SUPPLEMENTARY MATERIAL

#### FIGURES

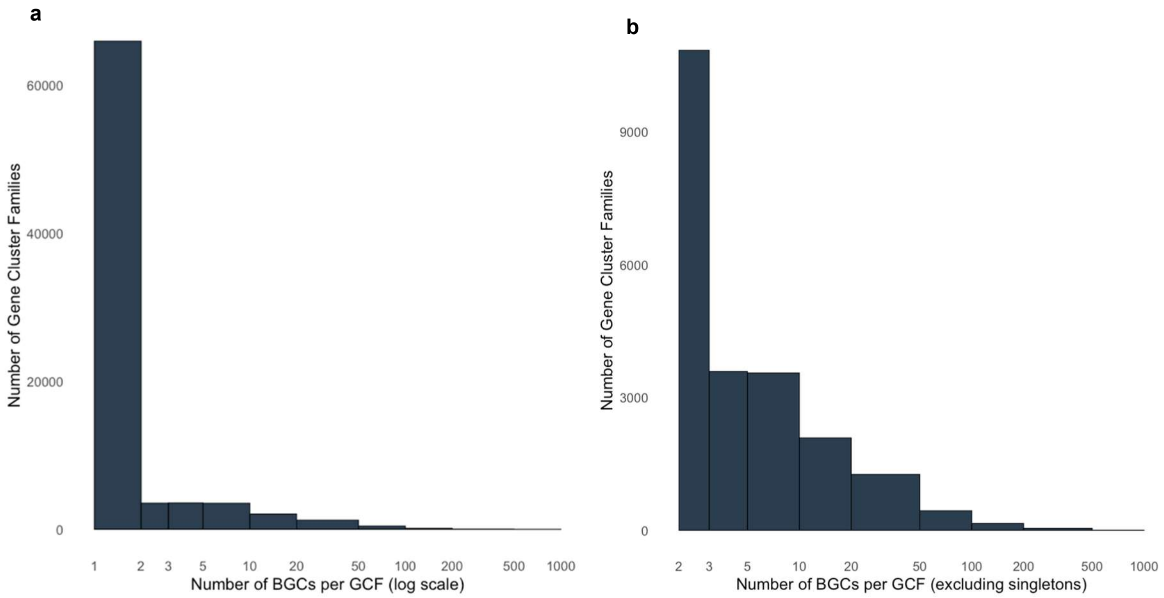


**Supplementary Figure 1:** Histograms showing the number of BGCs per GCF across the dataset. **a** GCF size distribution plotted on a logarithmic x-axis to highlight the long-tail structure of large families. **b** The same distribution shown on a linear scale, excluding singletons. The majority of GCFs consist of only a few BGCs, while a small subset include GCFs with hundreds of BGCs.

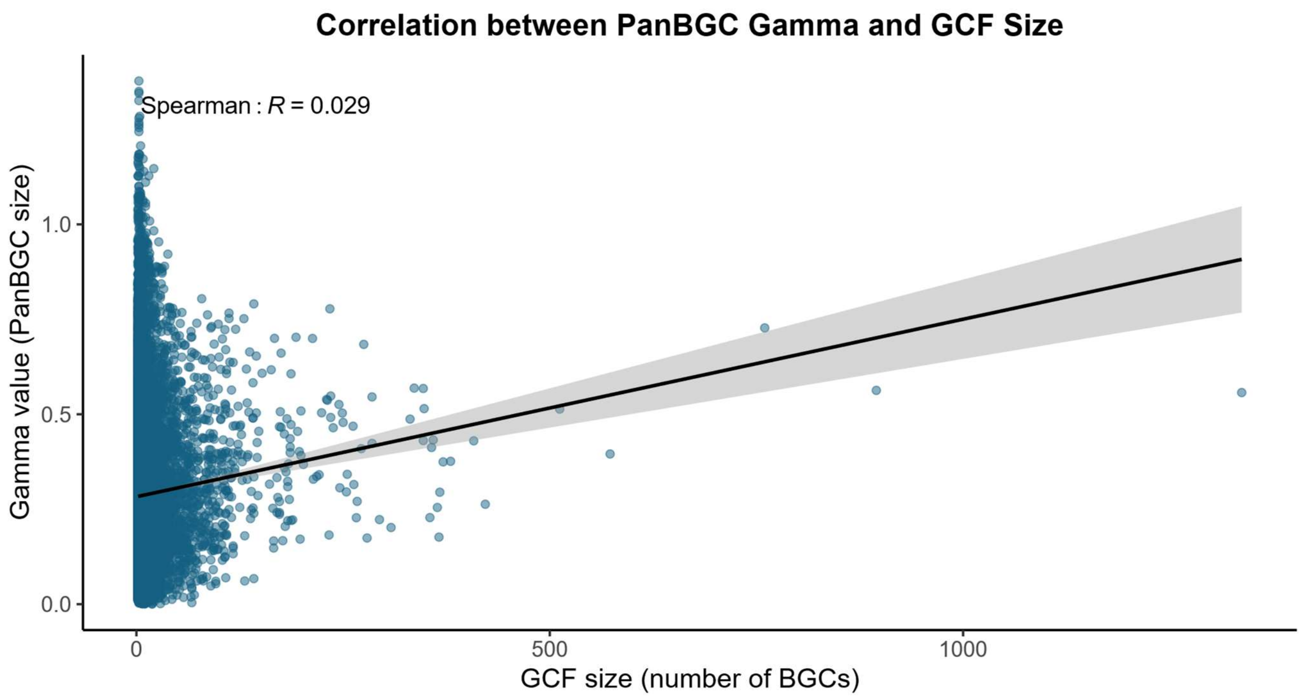


**Supplementary Figure 2: Correlation between GCF size and PanBGC openness (γ-value).** Scatterplot showing the relationship between the number of BGCs per gene cluster family (GCF size) and the corresponding gamma (γ) value calculated by the PanBGC framework, which quantifies openness based on Heaps' law. Each point represents a GCF. A slight positive trend is observed (Spearman's ρ = 0.029), indicating minimal correlation between family size and openness. The black line represents a linear regression fit with a 95% confidence interval (shaded area).

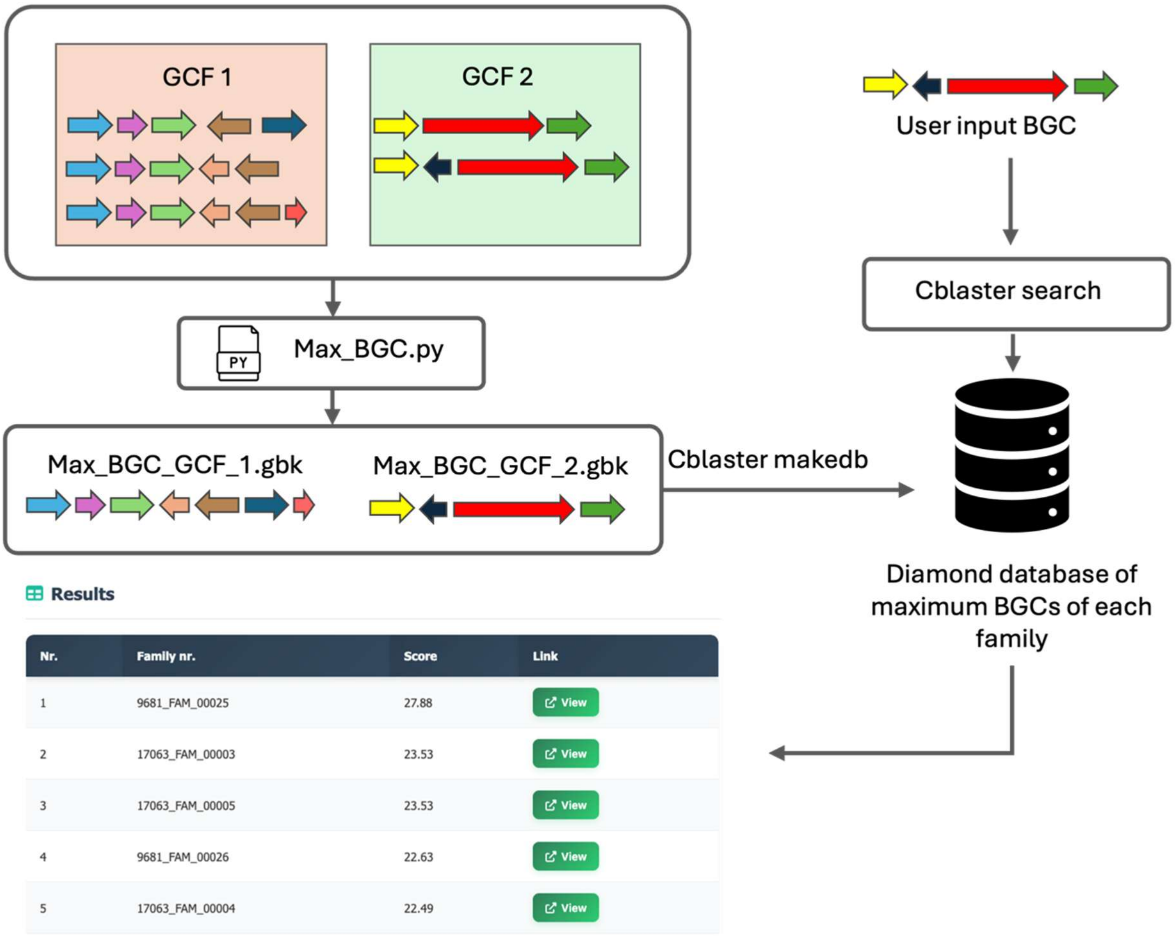


**Supplementary Figure 3: Cblaster database construction pipeline and user query.** Multiple GCFs results from ZOL are processed using a Python script (Max_BGC.py) to create a theoretical maximum BGC for each family. These maximum BGCs are then used to build a searchable DIAMOND database using the cblaster makedb module. A user-provided query BGC is subsequently searched against this database using cblaster search. The result identifies the best-matching family based on sequence similarity and hit coverage, which is displayed in a ranked table of candidate families.

#### TABLES

**Supplementary Table 1: Overview of json, excel and nexus files used for data storage.**

| File | Description |
|---|---|
| **Overview.json** | Contains summary information about all GCFs. Used for overview table creation. |
| **mibig_compound.json** | Contains information about mibig compounds and their family. Used for Compound overview table. |
| **BGC_analysis_results.xlsx** | Contains statistics of pfam domains found in each class. |
| **Gamma_value_bgc_data.json** | Contains summary information about the gamma calculation of each GCF. |
| **gbk_inf.json** | Available for each GCF. Contains information about each cluster in the GCF, and stores domain structure of each BGC |
| **genbank_data.json** | Available for each GCF. Contains different annotations for each gene and cluster part of the GCF |
| **Heaps_law.json** | Available for some GCF. Contains simulation order for heap's law calculation |
| **Nexus.nex** | Available for some GCF. Stores OG trees and the coalescent tree in nexus format. Used for tanglegram creation. |
| **Report.json** | Available for each GCF. Contains summary of ZOL run for the GCF. |

The files can be found under: https://github.com/ZiemertLab/PanBGC-DB/tree/master/Website_code/public/data
